# Feeding effects on liver mitochondrial bioenergetics of *Boa constrictor* (Serpentes: Boidae)

**DOI:** 10.1101/2021.07.05.451168

**Authors:** Helena Rachel da Mota Araujo, Marina Rincon Sartori, Claudia D. C. Navarro, José Eduardo de Carvalho, André Luis da Cruz

**Author notes:** These authors contributed equally to this work.

## Abstract

Snakes are interesting examples of overcoming energy metabolism challenges as many species can endure long periods without feeding, and their eventual meals are of reasonably large sizes, thus exhibiting dual extreme adaptations. Consequently, metabolic rate increases considerably to attend to the energetic demand of digestion, absorption and, protein synthesis. These animals should be adapted to transition from these two opposite states of energy fairly quickly, and therefore we investigated mitochondrial function plasticity in these states. Herein we compared liver mitochondrial bioenergetics of the boid snake *Boa constrictor* during fasting and after meal intake. We fasted the snakes for 60 days, then we fed a subgroup with 30% of their body size and evaluated their maximum postprandial response. We measured liver respiration rates from permeabilized tissue and isolated mitochondria, and from isolated mitochondria, we also measured Ca^2+^ retention capacity, the release of H_2_O_2_, and NAD(P) redox state. Mitochondrial respiration rates were maximized after feeding, reaching until 60% increase from fasting levels when energized with complex I-linked substrates. Interestingly, fasting and fed snakes exhibited similar respiratory control ratios and citrate synthase activity. Furthermore, we found no differences in Ca^2+^ retention capacity, indicating no increase in susceptibility to mitochondrial permeability transition pore (PTP), or redox state of NAD(P), although fed animals exhibited increases in the release of H_2_O_2_. Thus, we conclude that liver mitochondria from *B. constrictor* snakes increase the maintenance costs during the postprandial period and quickly improve the mitochondrial bioenergetics capacity without compromising the redox balance.

## INTRODUCTION

Mitochondria are complex and dynamic organelles present in eukaryotic cells responsible for energy production and cellular homeostasis. They play a fundamental role in the balance of energetic homeostasis upon intracellular signaling, apoptosis, metabolism of amino acids, lipids, cholesterol, steroids, and nucleotides, and its primary known function of oxidation of energetic substrates and ATP production (Duchen, 2000). This energy expenditure at the cellular level needs to be finely tuned to the varying availability of energy substrates from food resources and energetic demand from activities to allow better organismal performance. Animals can face challenges due to environmental changes as seasonal scarcity of food, behavior or life-history traits, increasing energy expenditure, like reproduction and migration. One basic regulation of energy expenditure depends on the control of oxidative phosphorylation, as this process accounts for most of the whole-animal oxygen consumption and has a considerable effect on cellular respiration flux (Benard et al., 2006; Brown et al., 1990; Dejean et al., 2001; Rolfe and Brown, 1997).

Ambush-foraging snakes are commonly used as experimental model organisms because of their resistance to long periods of food deprivation and the magnitude of their physiological responses after feeding large meals (Secor and Diamond, 1998; Starck and Beese, 2001; McCue et al., 2012). These snakes survive exceptional long periods of fasting by employing different strategies for energy conservation, as reduction in metabolic rates, organ mass and activity, and control of the mobilization of fuel sources (McCue, 2007; McCue et al., 2012). On the other hand, once fed, ambush-foraging snakes exhibit a remarkable increase in the metabolism, of comparatively higher magnitude than other animals (Secor and Diamond, 1998). The postprandial metabolic increment after meal intake (termed Specific Dynamic Action or SDA; Kleiber, 1961) may last for several days, depending on temperature regime, meal size, and quality (Andrade et al., 2004; Cruz-Neto et al., 1999; Gavira and Andrade, 2013; Secor and Diamond, 1997). Such elevated metabolism after feeding is mostly, if not fully fueled by aerobic metabolism. Thus, studies of the modulation of energy pathways involving oxidation of substrates ultimately leading to consumption of oxygen and production of ATP through the mitochondrial respiratory chain are essential to understand the regulation of metabolism at a cellular level.

In endothermic vertebrates, research has mainly focused on the mitochondrial effects of fasting, and studies conducted in mammals and birds report that food deprivation is accompanied by decreased mitochondrial respiration rates and increased rates of reactive oxygen species (ROS) production (Bourguignon et al., 2017; Dumas et al., 2004; Menezes-Filho et al., 2019; Roussel et al., 2019; Sorensen et al., 2006). Mitochondria unwittingly generate ROS as a by-product, and at low levels serves as redox signaling molecules, allowing adaptation to changes in environmental nutrients and oxidative environment (Schieber and Chandel, 2014; Shadel and Horvath, 2015). However, an excess can exhaust the antioxidant system and promote damage to proteins, lipids, and DNA, leading to oxidative stress (Hamanaka and Chandel, 2010). However, in species adapted to prolonged fasting, including mammalian hibernators, there seem to be mechanisms that allow the mitigation of oxidative stress (Ensminger et al., 2021). Nevertheless, although a robust body of literature exists for the physiological effects of fasting and feeding in snakes (McCue, 2008; Secor, 2009), knowledge of the optimization of metabolism at the subcellular level during periods of fasting or during the metabolic increment after meal intake are lacking (Butler et al., 2016).

We hypothesize that snakes will display mitochondrial plasticity, exhibiting an increase in the capacity for ATP generation during the postprandial period following the increase in energetic demand of digestion and absorption. To test this, we investigated the liver mitochondrial function and redox balance after 60-days of fasting and during the postprandial period in the ambush-foraging boid snake *Boa constrictor*. This neotropical snake feed infrequently, surviving periods of fasting longer than two months (McCue and Pollock, 2008), that can ingest large meals, exhibiting large increments in aerobic metabolic rate (Andrade et al., 2004; de Figueiredo et al., 2020; Toledo et al., 2003). As the liver plays a vital role in snake’s metabolism, participating in the oxidation of triglycerides, synthesis of cholesterol, lipoprotein and aminoacids, and control of blood sugar levels, it is relevant to assess the contribution of this organ to overall energetic demand after meal intake. In boas, the liver exhibit increased mass (Secor and Diamond, 2000) and a larger volume of glycogen granules two days post-feeding (da Mota Araujo, unpublished data). Thus, we compared mitochondrial liver bioenergetics of fasted and fed *B. constrictor*, evaluating mitochondrial respiration, calcium retention capacity, ROS release, and NAD(P) redox state.

## MATERIAL AND METHODS

### Reagents

We purchased the fluorescent probes Calcium Green™-5N and Amplex™ UltraRed from Thermo Fischer Scientific (Eugene, OR, USA) and dissolved in deionized water and dimethyl sulfoxide (DMSO), respectively. All other chemicals were obtained from Sigma-Aldrich (Saint Louis, MO, USA). Stock solutions of respiratory substrates and nucleotides were prepared in a 20 mM HEPES solution with the pH adjusted to 7.2 using KOH.

### Animals

We obtained juvenile snakes *Boa constrictor* Linnaeus, 1758 (N = 9, body mass = 152.09 ± 15.96; total length = 73.72 ± 3.35 cm, mean ± s.d.) from Centro de Recuperação de Animais Silvestres do Parque Ecológico do Tietê (CRAS, São Paulo, SP, Brazil). We housed the animals in individual boxes (56.4 l × 38.5 w × 20.1 h cm) with venting holes in the lid, under natural light and temperature (25 ± 2°C, mean ± s.d.) with free access to water. Initially, we fed all animals with mice (*Mus musculus*) to standardize the beginning of the treatment (with the equivalent of 5% of their body masses). After, we kept all snakes in fasting for two months. Then, we divided the snakes into two groups: fasting (N = 5) and fed (N = 4). We fed the snakes of the ‘fed group’ with mice accounting for 30% of their body weight and euthanized them 2 days after prey ingestion, usually when the maximum VO_2_ (oxygen consumption) is achieved (peak SDA; Secor and Diamond, 2000; de Figueiredo et al., 2020). We performed all measurements at the Laboratory of Bioenergetics at Universidade Estadual de Campinas (UNICAMP), Campinas, SP, Brazil. We anesthetized the snakes with isoflurane and sectioned the medulla after cessation of reflexes. All experimental procedures were approved by the Local Committee for Ethics in Animal Experimentation (CEUA/UNICAMP: 5301-1/2019) and complied with the ARRIVE guidelines. The use of *Boa constrictor* was authorized by the Brazilian Institute for Environment (SISBIO; number 69655-1).

### Permeabilized liver tissue

We rapidly removed a portion of the liver and immersed in ice-cold BIOPS buffer (10 mM Ca-EGTA buffer [2.77 mM of CaK_2_EGTA C 7.23 mM of K_2_EGTA, free concentration of calcium 0.1 mM], 20 mM imidazole, 50 mM KC/ 4- morpholinoethanesulfonic acid, 0.5 mM dithiothreitol, 7 mM MgCl_2_, 5 mM ATP, 15 mM phosphocreatine, pH 7.1). Then we permeabilized liver samples of 8 to 10 mg tissue in ice-cold buffer containing saponin (0.5 mg · mL^-1^) during 30 min, gently stirred and washed with MIR05 medium (60 mM potassium lactobionate, 1 mM MgCl_2_, 20 mM taurine, 10 mM KH_2_PO_4_, 20 mM HEPES, 110 mM sucrose, 1 g · L^-1^ BSA, pH 7.1) at 4°C. We dried the samples with filter paper and weighted (Busanello et al., 2017; Kuznetsov et al., 2008) before respirometric measurements.

### Mitochondrial isolation

We isolated liver mitochondria by conventional differential centrifugation (Ronchi et al., 2013). Briefly, we rapidly removed the liver, finely minced and homogenized in ice-cold isolation medium containing 250 mM sucrose, 1 mM EGTA, and 10 mM HEPES buffer (pH 7.2). We centrifuged the homogenate for 10 min at 800 *g*. Then, we centrifuged the collected supernatant at 7,750 *g* for 10 min. We resuspended the resulting pellet in buffer containing 250 mM sucrose, 0.3 mM EGTA, and 10 mM HEPES buffer (pH 7.2), and centrifuged again at 7,750 *g* for 10 min. We resuspended the final pellet containing liver mitochondria in an EGTA-free buffer at an approximate protein concentration of 60 mg · mL^-1^, quantified by the Bradford method using bovine serum albumin (BSA) as standards.

### Mitochondrial oxygen consumption

We measured mitochondrial respiration by monitoring the rates of oxygen consumption using a high-resolution oxygraph OROBOROS (Innsbruck, Austria), equipped with a magnetic stirrer, in a temperature-controlled chamber maintained at 30°C for permeabilized tissue and 28°C for isolated mitochondria. We suspended the permeabilized liver tissues in 2 mL of MIR-05 supplemented with 300 µM EGTA and 5 mM malate, 10 mM pyruvate, and 10 mM glutamate. After measuring the basal O_2_ consumption, respiration linked to oxidative phosphorylation (OXPHOS) was elicited by the addition of 400 µM of ADP. Then, we added 1 µg · mL^-1^ of oligomycin to cease the phosphorylation by ATP synthase (state 4_o_), slowing down oxygen consumption. Finally, we titrated carbonyl cyanide 4-(trifluoromethoxy) phenylhydrazone (FCCP) until maximal electron transport system capacity that occurred at the concentration of 800 ηM, eliciting maximal respiration rate (ETS, V_max_).

We suspended the isolated liver mitochondria (0.5 mg · mL^-1^) in 2 mL of standard reaction medium (125 mM sucrose, 65 mM KCl, 2 mM KH_2_PO_4_, 1 mM MgCl_2_, 10 mM HEPES buffer with the pH adjusted to 7.2 with KOH) supplemented with 200 µM EGTA and 1 mM malate, 2.5 mM pyruvate and 2.5 mM glutamate to evaluate respiration at complex I, with additions of 300 µM of ADP, 1 µg/mL of oligomycin and titration of FCCP, that elicited maximal respiration rate at 100 ηM. For isolated mitochondria we applied an additional protocol for the evaluation of the different mitochondrial complexes. We measured basal respiration with complex I-linked substrates (5 mM malate, 10 mM pyruvate and 10 mM glutamate), followed by the addition of ADP and FCCP as described above, then we added 1 µM rotenone to block complex-I followed by the addition of 5 mM succinate to stimulate complex II. Because the addition of 1 µM antimycin A or 1 µM myxothiazol were without effect on blocking complex III, we discarded the final addition of 1 mM *N,N,N*′,*N*′-tetramethyl-*p*-phenylenediamine (TMPD) plus 100 µM ascorbate aimed for stimulation of complex IV.

### Assessment of mitochondrial Ca^2+^ retention capacity

We suspended liver mitochondria (0.5 mg · mL^-1^) in standard reaction medium supplemented with 10 µM EGTA, 0.2 µM of a calcium indicator (Calcium Green™-5N) and respiratory substrates (1 mM malate, 2.5 mM pyruvate, and 2.5 mM glutamate). We continuously monitored the fluorescence in a spectrofluorometer (Hitachi F-4500, Tokyo, Japan) at 28°C using excitation and emission wavelengths of 506 and 532 nm, respectively, and slit widths of 5 nm. We performed repeated pulses of CaCl_2_ additions (60 µM) after mitochondria were added to the system. We measured the amount of CaCl_2_ added before the start of Ca^2+^ release by mitochondria into the medium as an index of the susceptibility to Ca^2+^-induced PTP, confirmed by the assessment of the PTP in the presence of 1 µM cyclosporine A (CsA). We converted the raw fluorescence readings into Ca^2+^ concentration levels (expressed as micromolar) according to the hyperbolic equation: [Ca^2+^] = K_d_ x [(F − F_min_)/(F_max_ − F)], where F is any given fluorescence, F_min_ is the lowest fluorescence reading after addition of 0.5 mM EGTA, and F_max_ is the maximal fluorescence obtained after two sequential additions of 1 mM CaCl_2_. We performed these additions of EGTA and Ca^2+^ at the end of each trace. We experimentally determined the dissociation constant (K_d_) of 26.8 µM for the probe Calcium Green™-5N in the incubation condition, as previously described (Sartori et al., 2021).

### Citrate synthase activity

We measured the catalytic activity of the enzyme citrate synthase in mitochondrial samples monitoring the conversion of oxaloacetate and acetyl-CoA to citrate and CoA– SH and by measuring the formation of the colorimetric product thionitrobenzoic acid (TNB) at 412 nm and 37°C (Shepherd and Garland, 1969) on a microplate reader (Power Wave XS-2, Biotek Instruments, Winooski, USA). We calculated the enzyme activity using the changes in absorbance after substrate (250 µM oxaloacetate) addition to the assay buffer (10 mM Trizma pH 8.0) containing 50 µM acetyl-CoA and 100 µM DTNB.

### Hydrogen Peroxide (H_2_O_2_) release

We monitored the H_2_O_2_ released by isolated liver mitochondria by the conversion of Amplex™ UltraRed to fluorescent resorufin in the presence of horseradish peroxidase (HRP). We incubated the suspensions of mitochondria from fasting and fed snakes (0.5 mg · mL^-1^) in a reaction medium containing complex-I substrates (1 mM malate, 2.5 mM pyruvate, and 2.5 mM glutamate), 10 µM Amplex™ UltraRed, 1 U · mL^-1^ HRP and 30 U · mL^-1^ superoxide dismutase (SOD). Additionally, we added 100 µM phenylmethyl sulfonyl fluoride (PMSF) to inhibit the conversion of amplex red by carboxylesterase independent of H_2_O_2_ (Miwa et al., 2016). We monitored the fluorescence over time with a temperature-controlled spectrofluorometer at 28°C (Hitachi F-4500, Tokyo, Japan) using excitation and emission wavelengths of 563 and 586 nm, respectively, and slit widths of 5 nm. For calibration, we added known amounts of H_2_O_2_ to the reaction medium with mitochondrial samples.

### NAD(P) redox state

We suspended the isolated liver mitochondria (0.5 mg · mL^-1^) in a standard reaction medium supplemented with 200 µM EGTA, and 5 mM succinate plus 1 µM rotenone, and monitored the changes in the redox state of NAD(P) in a spectrofluorometer (Hitachi F-7100) at 28°C, using excitation and emission wavelengths of 366 and 450 nm, respectively, and slit widths of 5 nm. Of note, only the reduced forms of NAD(P) exhibit a strong endogenous fluorescence signal. The peroxide-metabolizing system supported by NADPH was challenged with exogenous tert-butyl hydroperoxide (t-BOOH), an organic peroxide that is metabolized through the glutathione peroxidase/reductase system (Liu and Kehrer, 1996). As a reference, we added known amounts of NADH to the reaction medium in the absence of mitochondria.

### Statistical analyses

We tested for data normality and homoscedasticity by the Shapiro-Wilk and Barlett’s K-squared tests, respectively, using the R package. For variables that met the assumptions of parametric tests, we performed a two-tailed unpaired *t*-test for independent samples for comparison between fasted and fed. Whenever data failed the premises, we compared the groups by the Mann-Whitney test. We performed all analysis in Prism GraphPad software v. 7.1. We presented the results as mean and standard error (s.e.m.), assuming the significance level of 0.05.

## RESULTS

### Oxygen consumption of liver permeabilized tissue and isolated mitochondria

Liver permeabilized tissue from fed snakes exhibited 30% higher V_max_ than fasting snakes (**Fig. 1A**, unpaired *t*-test, *P*=0.0086). Citrate synthase activity of liver permeabilized tissue did not differ between the groups (**Fig. 1B**, unpaired *t*-test, *P*<0.5). For isolated mitochondria, fed snakes exhibited 40%, 58% and 64% higher respiration rates supported by complex I-linked substrates at basal, OXPHOS and, state 4_o_, respectively, in comparison to fasting snakes (**Fig. 2A, B**, *t*-test, *P*≤0.05). Mitochondrial V_max_ stimulated with complex I-linked substrates was 53% higher in the fed group (*t*-test, *P*≤0.05), while mitochondrial V_max_ stimulated with complex II-linked substrates were not different between fasting and fed (**Fig. 2C**, *t*-test, *P*=0.19). Because *B. constrictor* mitochondria were insensitive to both antimycin A and myxothiazol, we could not block complex III activity and could not evaluate complex III and IV. Mitochondrial respiratory control ratios and citrate synthase activity did not differ between groups (**Fig. 2D, E**, Mann-Whitney, *P*=0.45).

**Fig. 1.**
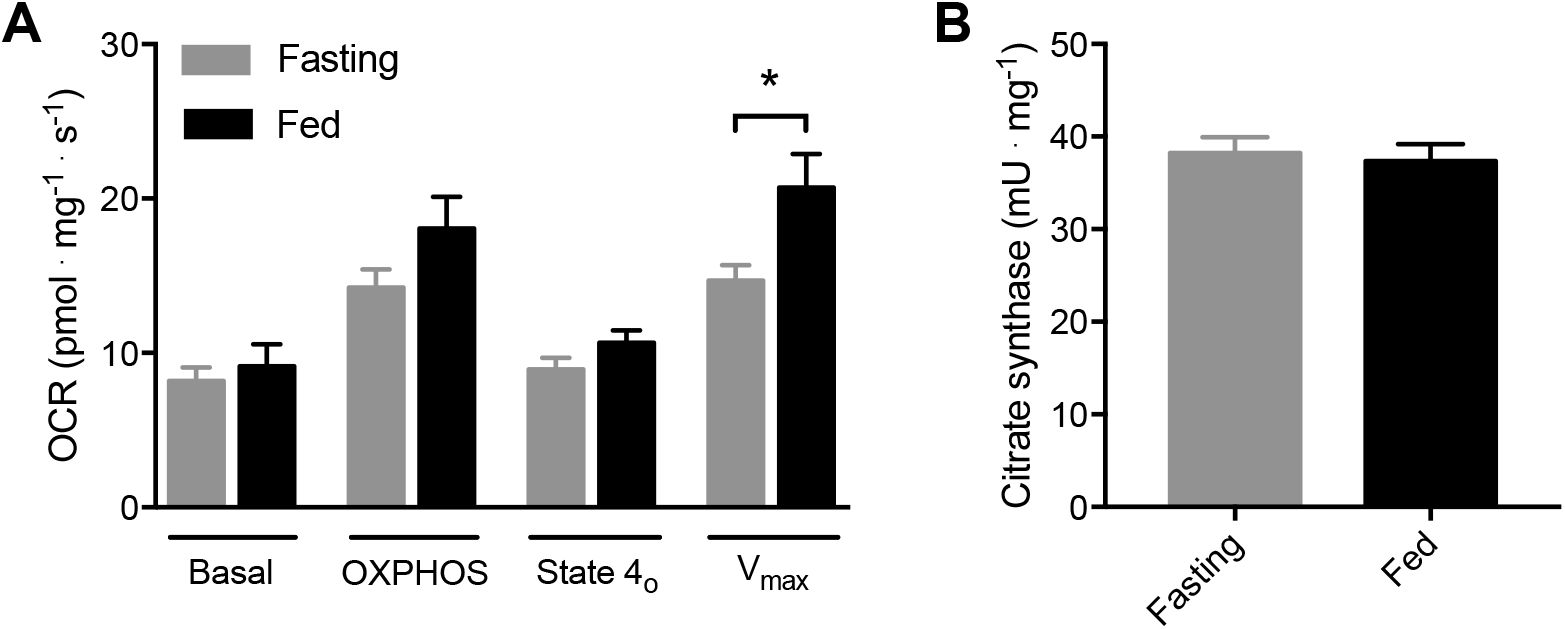
Feeding increased maximal respiration rate in liver permeabilized tissue from *B. constrictor* snakes. **A)** Oxygen consumption of liver permeabilized tissue at the presence of 5 mM malate, 10 mM pyruvate, and 10 mM glutamate as substrates (basal), after additions of 400 µM ADP (OXPHOS), 1 µg/ml oligomycin (State 4_o_), and 0.8 µM FCCP (V_max_). **B)** Citrate synthase activity. Bars denotes means ± s.e.m.; * *P*<0.01, *t*-test (N = 17 fasting, N = 11 fed; independent experiments).

**Fig. 2.**
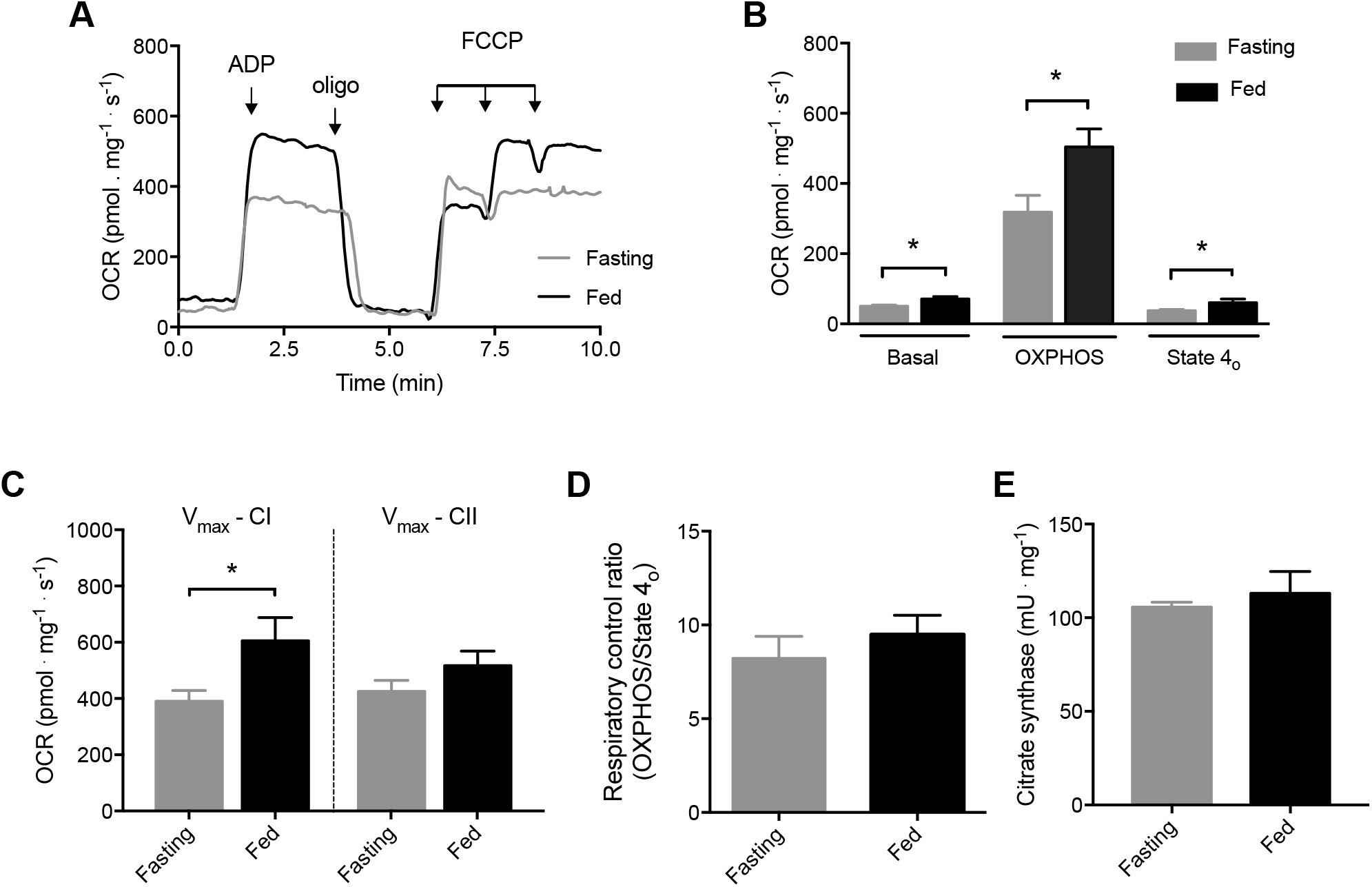
Feeding increased *B. constrictor* basal respiration, oxidative phosphorylation capacity, state 4_o_ and V_max_ in liver isolated mitochondria energized with complex I-linked substrates. **A)** Representative traces of oxygen consumption rate (OCR) of fasting and fed snakes, in the presence of 1 mM malate, 2.5 mM pyruvate and 2.5 mM glutamate as substrates, with additions of 300 µM ADP, 1 µg/ml oligomycin (oligo) and 50 ηM FCCP, where indicated by the arrows. **B)** Quantification of OCR per mg mitochondrial protein; \**P*≤0.5, *t*-test; **C)** Maximal respiration rate (V_max_) stimulated with complex-I and complex-II linked substrates; \**P*≤0.5, *t*-test; **D)** Respiratory control ratios (OXPHOS/State 4_o_); **E)** Citrate synthase activity. Data are presented as means ± s.e.m. (N = 5 fasting, N = 4 fed).

### Assessment of mitochondrial Ca^2^ retention capacity

Ca^2+^ retention capacity was evaluated by sequential additions of Ca^2+^ pulses (**Fig. 3A, B**) to the medium. Mitochondria of fasting and fed snakes exhibited similar capacities to retain calcium. Mitochondria of fasting snakes were able to take and retain approximately 264 ± 67 nmol Ca^2+^ · mg protein^-1^ versus 465 ± 79 nmol Ca^2+^ · mg protein^-1^ of fed snakes, (**Fig. 3C**, *t*-test, *P*>0.05). With the presence of CsA, both groups of snakes similarly increased resistance to PTP opening (1140 ± 35 nmol Ca^2+^ · mg protein^-1^ in the fasting group vs. 900 ± 173 nmol Ca^2+^ · mg protein^-1^ in the fed group) (**Fig. 3C**, Mann-Whitney, *P*=0.30).

**Fig. 3.**
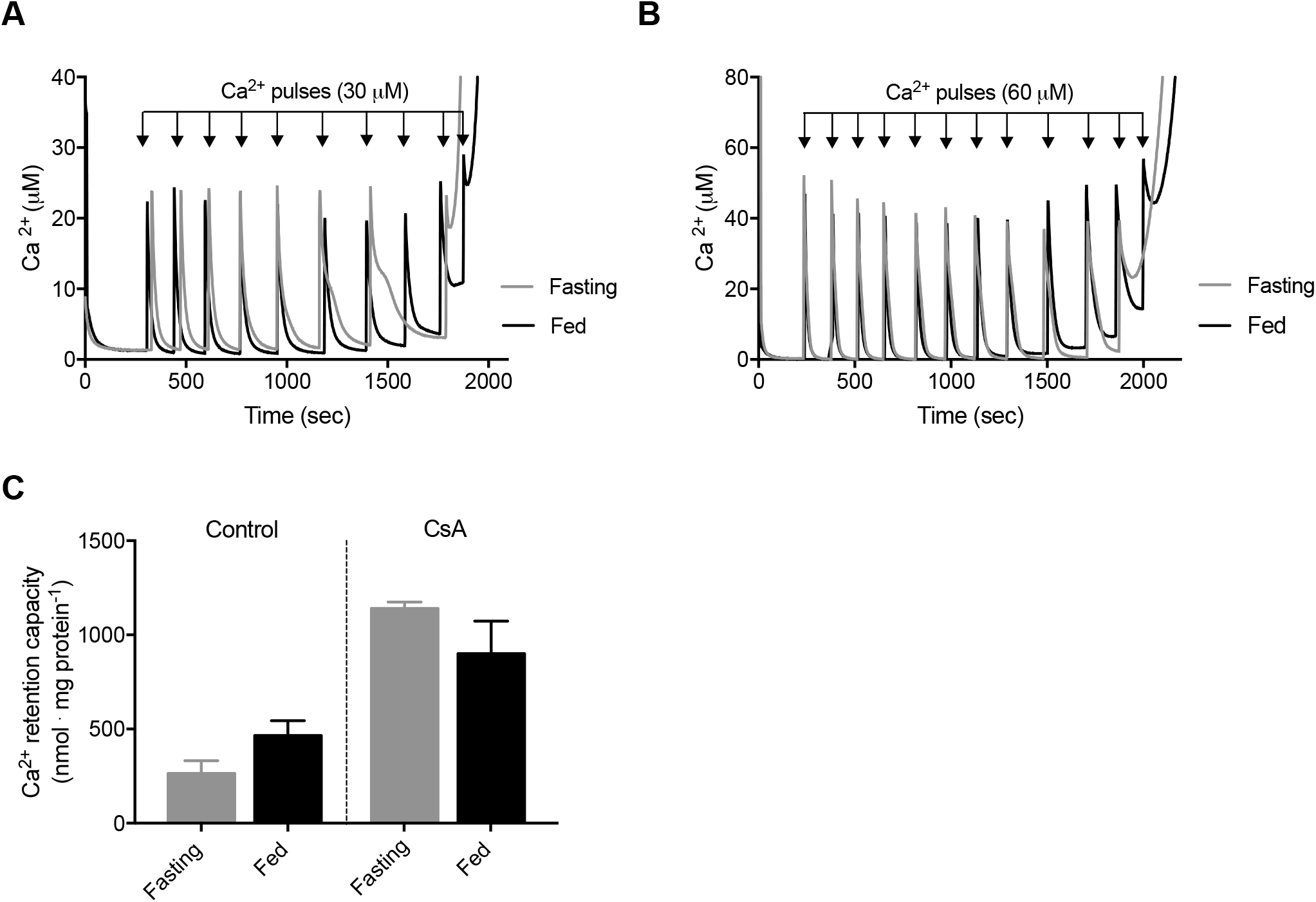
Feeding did not affect mitochondrial Ca^2+^ retention capacity in *B. constrictor* snakes. After addition of mitochondria to the system, Ca^2+^ retention capacity was accessed by consecutive additions of Ca^2+^ pulses until Ca^2+^-induced Ca^2+^ release as the consequence of the opening of PTP. Representative traces are depicted in **A)** control condition, Ca^2+^ pulses of 30 µM; **B)** in the presence of cyclosporine A (CsA), Ca^2+^ pulses of 60 µM. **C)** Amount of Ca^2+^ retained in each condition before the onset of permeability transition. Data are presented as means ± s.e.m., (N = 5 fasting, N = 4 fed).

#### Mitochondrial hydrogen peroxide (H_2_O_2_) release and redox state of NAD(P)

H_2_O_2_ released from liver mitochondria of fed snakes (76.5 ± 16.5 ρmol · mg^-1.^ min^-1^) was 2-fold higher than from fasting snakes (36.8 ± 2.5 ρmol · mg^-1.^ min^-1^) (Mann-Whitney, *P*=0.02) (**Fig. 4A, B**). There was no difference in the NADPH-dependent capacity to metabolize peroxide in fasting versus fed snakes (**Fig. 4C, D**, *t*-test, *P*=0.16).

**Fig 4.**
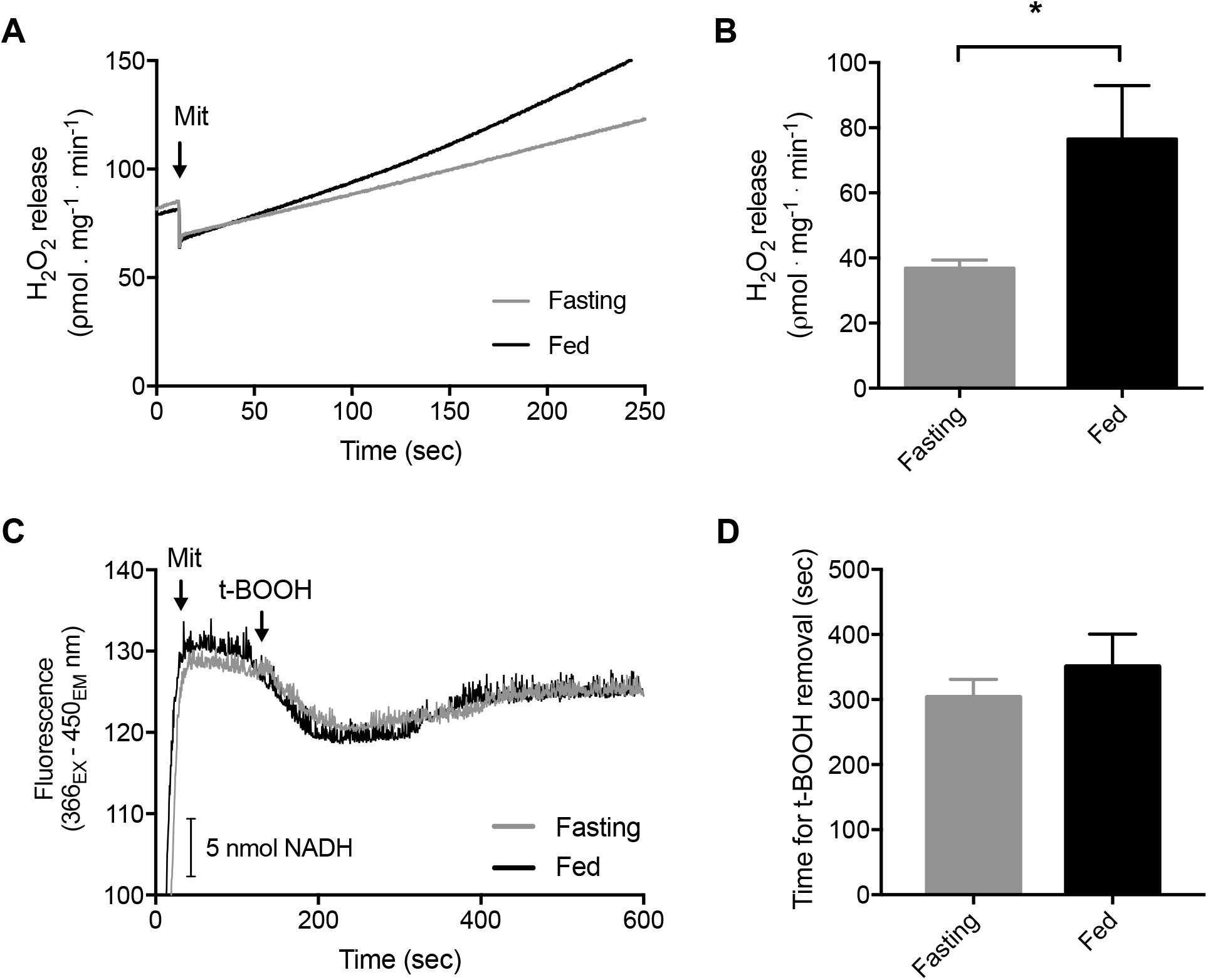
Liver mitochondria from fed individuals exhibited higher rates of H_2_O_2_ release but no evidence of oxidative stress in comparison to fasting individuals. **A)** Representative traces of H_2_O_2_ release assayed by the Amplex™ UltraRed probe in fasting (grey line) and fed (black line) liver mitochondria during resting respiratory state supported by complex I substrates. **B)** H_2_O_2_ released at a basal state (\**P*=0.03, Mann-Whitney test). **C)** Representative traces of the endogenous fluorescence of NAD(P) in the reduced state monitored over time in mitochondria incubated in standard reaction medium supplemented with succinate and rotenone in the absence of exogenous ADP; **D)** Time spent to recover the reduced state of NAD(P) following the addition of 2.5 µM organic peroxide (t-BOOH) load. The recovery time was used to indirectly estimate the rate of peroxide removal in each group and the ability of mitochondria to scavenge peroxide. Data are presented as means ± s.e.m. (N = 5 fasting, N = 4 fed).

## DISCUSSION

The present study revealed that *B. constrictor* liver mitochondria exhibit profound energetic changes in response to meal intake. After feeding, mitochondrial respiration rates from *B. constrictor* were significantly increased in comparison to unfed snakes. Mitochondria are dynamic structures, ongoing fusion, fission processes, and changes in number, morphology, and distribution, depending on the developmental, physiological, and environmental conditions (Mishra and Chan, 2016). Notwithstanding, the capacity to shift liver mitochondrial profiles two days after meal intake in boas is remarkable, bringing attention to the underlying mechanisms and the potential effects on mitochondria from other tissues directly or not involved in the digestion and absorption processes *per se*.

We observed a remarkable increase of liver respiration rate during OXPHOS (oxidative phosphorylation or state 3) of fed snakes compared to fasting, in the magnitude of approximately 60%. This increase reflects what is reported for whole-animal oxygen consumption rates (VO_2_) in snakes but should consider the differences in the meal size, the time spent at fasting, and the moment of post-feeding sampling. For example, varying periods of fasting in *B. constrictor* did not change the total energetic cost of digestion. However, it changed the temporal profile of the postprandial response (de Figueiredo et al., 2020). The increase in respiration rates seems to be fueled by substrates linked to complex I because we did not see differences in V_max_ between fasting and fed snakes when using complex II-linked substrates. Indeed, upregulation of genes for respiratory complex I, among other genes related to oxidative phosphorylation, was reported during digestion in snakes (Duan et al., 2017). Unfortunately, mitochondrial function studies in snakes are scarce. Interestingly, it was found that low temperature can impact coupling and efficiency in liver mitochondria of the snake *Natrix natrix*, but only when respiration was driven by succinate as the respiratory substrate (Dubinin et al., 2019), indicating that different sources of stimulus can impact mitochondrial function distinctly.

Another interesting finding was that feeding did not influence the quantity or efficiency of *B. constrictor* liver mitochondria since citrate synthase activity and respiratory control ratios were maintained. The mitochondria of fasting boas exhibited lower respiratory rates in all measured states, following the low resting energetic demand of the species (de Figueiredo et al., 2020; Stuginski et al., 2018). However, the capacity to also exhibit a lower respiration rate after ATP synthase blockade with oligomycin (state 4_o_) indicates the capacity to reduce leakage of protons through the membrane, which is a crucial contributing factor towards energy saving (Brand et al., 1993), and probably to the low metabolic rates observed in ambush-hunting snakes (Stuginski et al., 2018). Unfortunately, we could not determine the respiration induced by the proton leak due to the lack of response to complex III inhibition. The insensitivity to complex III inhibitors indicates that the ubiquinol-cytochrome c oxidoreductase complex may exhibit a different molecular structure in snakes, as this outcome was also observed in *Bothrops alternatus* (Ogo et al., 1993) and *Python regius* (Bundgaard et al., 2020). Like boas, long-term fasting in king penguin chicks also did not reduce skeletal muscle mitochondria efficiency compared to short-term fasting in birds (Bourguignon et al., 2017). In contrast, fasting mammals exhibited the same or even more significant proton leak than after feeding, compromising the efficiency of mitochondria during food deprivation periods (Brown and Staples, 2011; Menezes-Filho et al., 2019; Sorensen et al., 2006). Therefore, the capacity of lowering the leakiness might imply an adaptation of the *B. constrictor* mitochondria to recurrent fasting periods.

In contrast to findings from isolated mitochondria, maximal respiration rate (V_max_) of permeabilized liver tissue was the only difference between our fasting and fed boas. The different results obtained from liver isolated mitochondria and permeabilized tissue could reflect intracellular interactions (Picard et al., 2010) or relate to different bioenergetics profiles exhibited by mitochondrial subpopulations. Recently, a paper investigating brown adipose tissue (BAT) mitochondria showed that when associated with lipid droplets, mitochondria exhibited increased coupling, related to fatty acid synthesis, in contrast to cytoplasmic mitochondria, which were related to fatty acid oxidation (Benador et al., 2018). Interestingly, the bioenergetics results of the lipid droplet subpopulation of mitochondria from BAT of Benador’s work were similar to the results obtained from *B. constrictor* of the fed treatment. Also, snakes preferentially oxidize protein over lipids during the 14 days after feeding (McCue et al., 2015), increasing the proportion of lipid droplet mitochondria compared to cytoplasmic mitochondria in the liver during the postprandial period. Nevertheless, more work is needed to investigate if there are bioenergetics differences from mitochondrial subpopulations in boas.

The mitochondrial Ca^2+^ retention capacity is a proxy for evaluating susceptibility of the mitochondrial permeability transition pore (PTP), a phenomenon characterized by the Ca^2+^-dependent opening of a non-specific pore in the inner mitochondrial membrane. The PTP affects the structure and function of mitochondria, which is ultimately related to cell death by apoptosis or necrosis and to many pathological conditions (Vercesi et al., 2018). The amount of Ca^2+^ that leads to overload, thus triggering PTP, varies with the source and conditions of mitochondria and the presence of protectors or inducers acting on the still debated pore constitutional units (Kowaltowski et al., 2001). Differently from mice mitochondria, which showed a higher susceptibility to PTP when at fasting (Menezes-Filho et al., 2019), nutritional status does not seem to affect the susceptibility to PTP in *B. constrictor*, as fasting and fed snakes exhibited no differences in mitochondrial Ca^2+^ retention. PTP can be sensitized by oxidative stress and oxidized NADPH (NADP^+^) (Castilho et al., 1995; Vercesi et al., 1988; Zago et al., 2000), as excess ROS increase oxidation of protein thiols and promotes disulfide bonds and cross-linked protein aggregation in the inner mitochondrial membrane (Castilho et al., 1995; Fagian et al., 1990; Valle et al., 1993; Vercesi, 1984). However, as we will discuss further, we also did not observe changes in NAD(P) redox status, and the increased rate of H_2_O_2_ production seems not to be leading to oxidative stress, thus not influencing PTP sensibility.

Liver mitochondria from *B. constrictor* exhibited higher rates of H_2_O_2_ released after ingestion of a meal compared to fasting, which contrasts with mitochondria from fed mammals and a study in fasted fish. Rodent mitochondria and the brown trout (*Salmo trutta*) exhibited higher levels of released H_2_O_2_ when subjected to fasting (Menezes-Filho et al., 2019; Salin et al., 2018; Sorensen et al., 2006). The reduced H_2_O_2_ of fasting boas may be due to the low energetic demand during fasting in snakes (Ensminger et al., 2021) and may be related to the remarkable capacity of metabolic regulation in such animals (McCue, 2007). Nevertheless, a two-month period fasting in *B. constrictor* may not be sufficient time to induce detrimental effects in mitochondria. Ambush-hunting snakes were shown to possess lower metabolic rates than active foraging snakes that feed more frequently (Stuginski et al., 2018), meaning that the energetic costs could be sustained for long periods using stored energy reserves. For example, the rattlesnake *Crotaluss durissus* was shown to endure under 12 months of food deprivation with slow body mass loss and no changes in resting VO_2_ (Leite et al., 2014). The increase in H_2_O_2_ could simply reflect the increase in aerobic metabolism and seems to not directly lead to oxidative stress and damage because we observed no differences in the redox status of NAD(P). Also, there is evidence that increased ROS generation after feeding in snakes may be concurrent to increased antioxidant defense, as genes encoding antioxidant enzymes like catalase, peroxiredoxin, glutathionine transferase, and heat shock protein were shown to be upregulated in digesting pythons (Duan et al., 2017).

Studies are increasingly showing that ROS generation is not essentially connected to damage, with demonstrations that ROS can act as signaling molecules, playing an essential role in the crosstalk from mitochondria and nucleus to maintain cell homeostasis (Shadel and Horvath, 2015). Of note, in mammals, there are remarkable differences between an acute fasting event and chronic fasting regimes as intermittent fasting (IF) or caloric restriction (CR) interventions. In both IF and CR, there is growing evidence that chronic recurrent fasting regimes improve defenses against oxidative stress and repair of damaged molecules (de Cabo and Mattson, 2019). In liver mitochondria from rodents, caloric restriction did not affect respiration rates but reduced ROS generation when energized with complex I-linked substrates and protected against PTP (Lambert et al., 2004; López-Torres et al., 2002; Menezes-Filho et al., 2017). In *B. constrictor*, which is adapted to recurrent fasting regimes, similar adaptive mechanisms can be potentially operative. Nevertheless, more studies could be performed to carefully evaluate the contrasting effects of transient beneficial effects of ROS and harmful sustained elevated ROS levels in response to a fasting-feeding transition in snakes of different feeding strategies.

## CONCLUDING REMARKS

In summary, our results showed that liver mitochondria of *B. constrictor* possess postprandial effects, exhibiting a rapid shift of mitochondrial bioenergetics towards higher respiration rates and oxidative phosphorylation supported by complex I-linked substrates, demonstrating the plasticity of snakes’ mitochondrial function. Furthermore, our results showed that mitochondrial function adaptations of boas might play a vital role in the fasting and feeding transition and be pivotal in organismal fitness by affecting animal performance.

## Acknowledgements

The authors are thankful to Prof. Anibal E. Vercesi (University of Campinas, SP, Brazil) for providing all the necessary facilities to conduct this research work, and to “Centro de Recuperação de Animais Silvestres do Parque Ecológico do Tietê (CRAS, São Paulo, SP, Brazil)”, for the provision of animals.

## Author contributions

Conceptualization: ALC, HRMA, MRS; Methodology: HRMA, MRS, CDCN; Formal analysis: HRMA, MRS; Investigation: HRMA, MRS, CDCN; Resources: MRS, CDCN, JEC; Writing - original draft: HRMA, MRS; Writing - review & editing: HRMA, MRS, ALC, JEC; Supervision: ALC, JEC; Project administration: ALC, JEC; Funding acquisition: ALC.

